# Adaptive functions correlate with evoked neurotransmitter release in SYT1-associated neurodevelopmental disorder

**DOI:** 10.1101/2023.11.25.568507

**Authors:** Paul Y. Park, Lauren E. Bleakley, Nadia Saraya, Reem Al-Jawahiri, Josefine Eck, Marc A. Aloi, Holly Melland, Kate Baker, Sarah L. Gordon

## Abstract

Pathogenic missense variants in the essential synaptic vesicle protein synaptotagmin-1 (SYT1) cause a neurodevelopmental disorder that is characterised by motor delay and intellectual disability, hyperkinetic movement disorder, episodic agitation, and visual impairments. SYT1 is the presynaptic calcium sensor that both triggers and drives synchronous neurotransmitter release. We have previously shown that pathogenic variants around the calcium-sensing region of the critical C2B domain decrease synaptic vesicle exocytosis in neurons. Here we show that recently identified variants within the facilitatory C2A domain of the protein (L159R, T196K, E209K, E219Q), as well as additional variants in the C2B domain (M303V, S309P, Y365C, G369D), share this underlying pathogenic mechanism, causing a graded and variant-dependent dominant-negative impairment in exocytosis. We establish that the extent of disruption to exocytosis *in vitro* correlates with neurodevelopmental impacts of this disorder. Specifically, the severity of motor and communication impairments exhibited by individuals harbouring these variants correlates with multiple measures of exocytic impairment. Together, this suggests that there is a genotype-function-phenotype relationship in SYT1-associated neurodevelopmental disorder, centring impaired evoked neurotransmitter release as a common pathogenic driver of this disorder. Moreover, this points toward a direct link between control of neurotransmitter release and development of adaptive functions, and provides a tractable target for therapeutic amelioration.

## Introduction

High-fidelity neurotransmission is achieved through tight coupling of action potentials at nerve terminals to synaptic vesicle exocytosis. In the cerebrum, this is primarily mediated by the presynaptic vesicular calcium sensor synaptotagmin-1 (SYT1) (Melland et al., 2021). SYT1 inhibits vesicle fusion in resting neurons (Yoshihara & Littleton, 2002; Xu et al., 2009); upon the influx of calcium through voltage-gated calcium channels, calcium binds to SYT1, acting as an electrostatic switch allowing the protein to penetrate the plasma membrane (Shao et al., 1997). This triggers and facilitates the fusion of neurotransmitter-laden synaptic vesicles with the presynaptic plasma membrane (Bai et al., 2004), and thus the release of neurotransmitters into the synaptic cleft. Complete loss of SYT1 results in early lethality and a profound abrogation of evoked neurotransmitter release in excitatory and inhibitory neurons with a concomitant increase in both spontaneous and asynchronous release (Geppert et al., 1994; Yoshihara & Littleton, 2002).

The two calcium-binding C2A and C2B domains of SYT1 cooperate in membrane binding and vesicle fusion. Calcium binding to the C2B domain is essential for triggering synchronous evoked release (Lee et al., 2013; Xu et al., 2009). This domain interacts with the pre-fusion SNARE complex to effectively position SYT1 to respond to calcium influx, and is an energetic driver of membrane fusion (Wang et al., 2016; Gruget et al., 2020). In comparison, the C2A domain plays an important facilitatory role in vesicle fusion by helping activate the electrostatic switch that triggers synchronous release (Striegel et al., 2012), while also demonstrating functional cooperativity with the C2B domain (Lee et al., 2013; Gruget et al., 2020). The lack of a functional C2A domain results in a significant reduction, though not complete ablation, of evoked neurotransmitter release (Paddock et al., 2011; Lee et al., 2013; Bowers & Reist, 2020; Wu et al., 2022). Dysfunctional SYT1, caused by variants in either C2 domain, perturbs both excitatory and inhibitory neurotransmission (Shields et al., 2020; Shin et al., 2009; Zhou et al., 2017; Wu et al., 2017). Furthermore, disruption to normal SYT1 function in mammalian systems can significantly impair both short-term (Bouazza-Arostegui et al., 2022) and long-term (Wu et al., 2017) plasticity, as well as altering short-term depression (Fernandez-Chacon et al., 2002). This highlights the relevance of SYT1 and neurotransmission kinetics for the synaptic correlates of learning and cognition. However, studies investigating behavioural impacts of SYT1 variation in mammalian model systems remain limited (Geppert et al., 1994; Powell et al., 2004).

Heterozygous *de novo* missense variants in *SYT1* cause a rare condition known as SYT1-associated neurodevelopmental disorder (OMIM #618218, Baker-Gordon Syndrome or BAGOS) (Baker et al., 2015; Baker et al., 2018). Frequently reported clinical features of this disorder include infantile hypotonia, congenital ophthalmic abnormalities, early -onset hyperkinetic movement disorders, and developmental delay / intellectual disabilities (Baker et al., 2015; Baker et al., 2018; Melland et al., 2022). A characteristic feature of the condition is electroencephalogram (EEG) abnormalities (most often low frequency, high amplitude oscillations), which is not accompanied by overt seizures or abnormal structural neuroimaging in most cases (Baker et al., 2015; Baker et al., 2018).

The first human pathogenic variants identified in SYT1 consisted of single amino acid substitutions in residues within the Ca^2+^-binding loops of the C2B domain of the protein (Baker et al., 2015; Baker et al., 2018). *In vitro* studies revealed that, while one variant (M303K) caused a reduction in protein stability and expression, all other variants (D304G, D366E, I368T and N371K) induced dominant-negative impairment of synaptic vesicle exocytosis (Baker et al., 2015; Baker et al., 2018; Bradberry et al., 2020). These findings provided the first evidence of a pathogenic mechanism underlying the condition. This disruption to exocytosis was demonstrated to be reversible by increasing intracellular calcium concentrations, either via application of increased extracellular calcium or the potassium-channel blocker 4-AP (Baker et al., 2018; Bradberry et al., 2020).

Recently, additional disorder-associated SYT1 variants have been identified, both within the C2B domain (including outside of the calcium-binding pocket) and within the C2A domain of the protein (Melland et al., 2022; Huang et al., 2023; Cornelisse et al., 2023). Molecular dynamics simulations demonstrated a lack of convincing evidence of a major structural component to the mechanisms of pathogenicity for most of these variants (Melland et al., 2022), and a pathogenic mechanism for these novel variants is yet to be established. In parallel with the increasing number and diversity of reported SYT1 variants, standardised phenotyping revealed a broadened range in the severity of associated neurodevelopmental characteristics (Melland et al., 2022). The extent of developmental delay and intellectual disability varied from mild to profound; 66% of 22 patients were reported to have a movement disorder, ranging from isolated ataxia or dystonia to severe mixed hyperkinetic involuntary symptoms (Melland et al., 2022). Other common features included visual impairments, emotional instability and self-injurious behaviours, which occurred at higher frequency than observed in a comparison group with other monogenic neurodevelopmental disorders, but varied widely in severity within the SYT1 group. A simple dichotomy in the severity of symptoms could not be found between individuals with C2A or C2B domain variants, suggesting that a more complex genotype-specific impact on SYT1 function might influence phenotypic severity.

Building on these previous observations, the current study had two major objectives. Firstly, we aimed to identify impairments to neuronal physiology induced by novel variants in the C2A and C2B domains of SYT1, in order to establish their pathogenicity and identify potential diversity of functional impacts. We specifically investigated missense variants across the two domains that were predicted to have differential impacts on SYT1 structure and function (Melland et al., 2022). Secondly, we interrogated whether the observed functional impacts contribute to variable severity of the associated neurodevelopmental condition. The recurrent C2B variant I368T has well-characterised functional impacts, and hence was included as a reference variant (Baker et al., 2015; Baker et al., 2018; Melland et al., 2023). We reveal here a shared mechanism of pathogenicity that correlates with core neurodevelopmental features of SYT1-associated neurodevelopmental disorder.

## Materials and Methods

### Functional studies

Full detail of materials and all methods can be found in the Supplementary Material.

All experiments were performed in accordance with the Prevention of Cruelty to Animals Act, 1986 under the guidelines of the National Health and Medical Research Council (NHMRC) of Australia Code of Practice for the Care and Use of Animals for Experimental Purposes. All experiments were approved by the Animal Ethics Committee at the Florey Institute of Neuroscience and Mental Health prior to commencement.

Site-directed mutagenesis was used to introduce the human SYT1 variants into homologous positions in rat SYT1 (human/rat: L159/158>R, T196/195>K, E209/208>K, E219/218>Q, M303/302>V, S309/S308>P, Y365/364>C, G369/368>D, I368/367>T, amino acid numbering used henceforth follows the human sequence), that was fused at the N-terminal lumenal domain to a pH-sensitive EGFP (superecliptic pHluorin (Sankaranarayanan et al., 2000)). Mutagenesis was performed using the QuikChange II Site-directed Mutagenesis kit (Agilent Technologies).

Primary hippocampal-enriched neuronal cultures were prepared from embryonic day 16.5–18.5 C57BL/6J mouse embryos. Cells were co-transfected with SYT1-pHluorin variants and either mCherry or empty vector pcDNA3.1 after 6-8 days and either fixed or imaged live after 13-15 days in culture.

For SYT1 expression analysis, cultured hippocampal cells were fixed, immunolabelled for SYT1 and GFP and subsequently imaged with a Zeiss Axio Observer 7 inverted epifluorescence microscope with a Colibri 7 LED light source. Images were processed offline using ImageJ 1.51s software, where circular 8x8 pixel regions of interest (ROIs) were chosen over nerve terminals to specifically measure fluorescence intensities within these regions (see Supplementary Methods for further details).

For live imaging assays, cultured hippocampal neurons harbouring SYT1-pHluorin variants were mounted in a Warner imaging chamber with embedded parallel platinum wires (RC-21BRFS) before being placed on the stage of a Zeiss Axio Observer 7 inverted epifluorescence microscope for imaging. To assess membrane partitioning of SYT1 variants, neurons were sequentially perfused with saline buffer, acidic MES buffer, or NH_4_Cl buffer. To assess impacts on exocytosis, neurons were perfused with saline buffer supplemented with 10 µM CNQX, 50 µM DL-AP5 and 1 µM vATPase inhibitor bafilomycin A1 at 37°C and stimulated with a train of 1200 action potentials at 10Hz.

### Participant recruitment and phenotyping

*SYT1* variants were identified via exome sequencing (trio or solo) within clinical laboratories or ethically approved research studies as previously reported (Melland et al., 2022; Baker et al., 2018). The identification, validation, confirmation of *de novo* status and clinical reporting of *SYT1* variants were carried out by each participant’s clinical centre. Authors were notified of diagnosed variants by personal communication, through database searching of ClinVar (Landrum et al., 2018), Decipher (Firth et al., 2009), or through Gene-Matcher (Sobreira et al., 2015). Written informed consent for inclusion in the current study was provided by each diagnosed individual’s parent or consultee (Cambridge Central Research Ethics Committee ref IRAS 83633: “Phenotypes in Intellectual Disability”). Clinical information was collated from medical documentation. Parent-report questionnaires were completed online or by post or by online interview as reported (Melland et al., 2022). The study-specific Medical History Interview gathered information about perinatal history, infant and child health, neurological symptoms, and developmental milestones. The number of reported movement disorder types was summed into an ordinal score. The Vineland Adaptive Behaviour Scales (third edition)(Sparrow & Cicchetti, 2016) is a standardized assessment tool for everyday adaptive functioning domains (communication, daily living skills, socialisation, and motor skills). The Developmental Behaviour Checklist 2 (DBC)(Gray et al., 2018) assesses emotional and behavioural problems in individuals with neurodevelopmental disorders and comprises 5 subscales (disruptive, self-absorbed, communication disturbance, anxiety and social relating). In line with our previous work, analysis focused on five items reflecting frequently-reported symptoms within the *SYT1* group (6. Bangs head; 10. Chews or mouths body parts; 33. Hits or bites self; 47. Mood changes rapidly for no reason; 60. Has repeated movements of hands, body or head). These items appear across two DBC-P subscales (disruptive/antisocial, self-absorbed). Raw scores (0-2) for each item were summed for each participant. The Cerebral Visual Impairment (CVI) scale (Ortibus et al., 2011) comprises six subscales (visual attitude, ventral stream, dorsal stream, complex visuomotor abilities, other senses, and associated characteristics), and analysis focused here on total scores. *SYT1* group data is outlined in Supp Table 2 and 3.

### Statistical analyses

Experimenters were blinded to the SYT1-pHluorin variant for image acquisition and analysis for fixed assays, and for image analysis for live imaging. Statistical analyses of SYT1 variant functional data were performed using GraphPad Prism Version 9.4.0 and Microsoft® Excel software. Each data set was tested for normality using the Shapiro-Wilk test and statistical analyses were performed accordingly (one-way ANOVA with Dunnett’s multiple comparison test for data with normal distribution, non-parametric Kruskal-Wallis test with Dunn’s multiple comparisons for data that did not conform to normal distribution, with each SYT1 variant protein compared to the wild-type (WT) protein). Time courses of change in fluorescence were analysed via repeated measures mixed model ANOVA with Dunnett’s multiple comparison test, to compare each SYT1 variant protein to the WT protein at every individual time point. For all analyses, p < 0.05 was considered statistically significant. ‘n’ refers to an individual field of view from an independent coverslip. All experiments were repeated across at least 3 independent cultures, with each culture comprising at least 3 embryos.

Permutation-based Spearman’s correlation tests were performed to examine associations between functional activity (measures of *in vitro* synaptic vesicle exocytosis kinetics) and quantitative phenotyping measures for participants with the equivalent SYT1 variants. The SYT1 variants included in this analysis, based on available functional and phenotypic data, are the following: T196K, E209K, E219Q, M303V, S309P (DBC data not available), Y365C, G369D (all variants n = 1 participants), and I368T (n = 6 participants, mean scores assessed). To account for multiple comparisons, the false discovery rate was controlled using the Benjamini-Hochberg procedure, with q value < 0.05 (p ≤ 0.019). Correlations with CVI total score and DBC self-injury items score controlled for age. For other measures, bivariate correlation tests were appropriate as the variables represented standardised scores accounting for age.

## Results

### Most SYT1 variants do not impact SYT1 expression or trafficking

We first explored whether newly-identified SYT1 variants (Fig 1a) impact the expression of the protein in neurons (Fig 1b). When transfected into neuronal cultures, the majority of SYT1 variants were expressed at a similar level to the control WT protein (Fig 1c), which suggests that protein stability is not compromised by most variants. Additionally, SYT1 levels in transfected cells (containing both exogenous protein and endogenous wild-type protein) were approximately twice as high as that observed in untransfected cells (with endogenous wild-type protein only) for most variants. This equates to an approximately equal copy number of exogenously introduced variant and endogenous wild-type protein, thereby mimicking the heterozygous expression of these variants. However, the L159R variant is a notable exception, as it was expressed at a significantly lower level than the WT protein (Fig 1c). The L159R variant introduces a charged, polar arginine that faces into the hydrophobic interior of the C2A domain, which molecular dynamics simulations predicted may cause notable, diffuse structural perturbations within the domain (Melland et al., 2022). Together, these lines of evidence indicate that L159R likely disrupts SYT1 stability.

**Figure 1.**
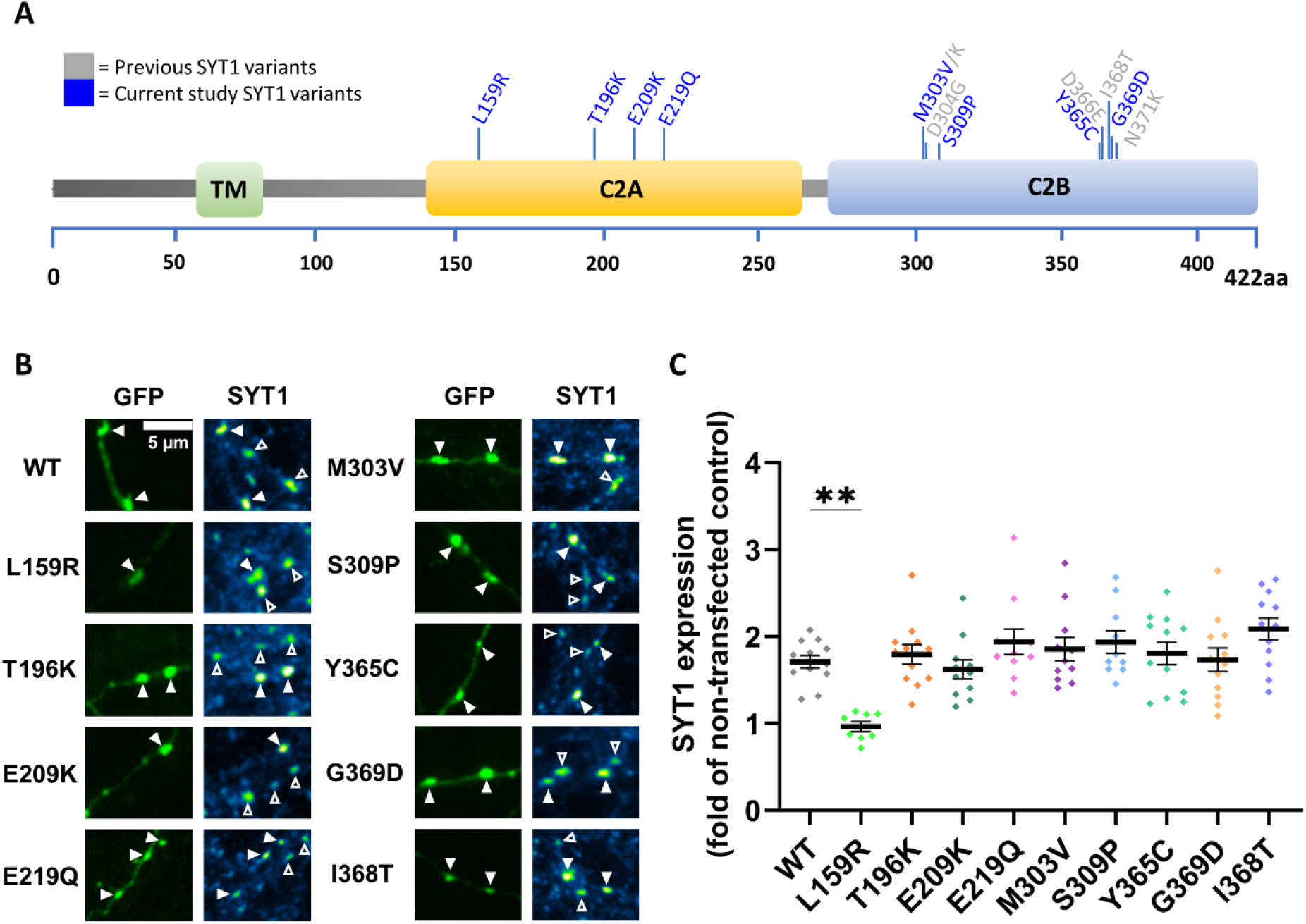
SYT1 variants, except L159R, are expressed as efficiently as the WT protein at presynaptic nerve terminals. Hippocampal neurons transfected with SYT1-pHluorin variants were fixed and immunolabelled for GFP and SYT1. (**A**) Linear protein sequence of SYT1 highlighting the location of previously investigated (grey) and currently examined (blue) SYT1 variants. (**B**) Representative images of immunolabelled neurons transfected with SYT1 variants (tagged with pHluorin). Left panels show GFP-transfected neurons in green and right panels display SYT1 immunofluorescence, with warmer colours indicating more intense staining. Arrowheads indicate transfected (closed) and non-transfected (open) nerve terminals. Scale bar = 5µm. (**C**) SYT1 expression levels, expressed as the SYT1 immunofluorescence intensity in transfected neurons, relative to non-transfected neurons in the same field of view. Data displayed as mean ± SEM, n = 8-12 (from 3 independent cultures). Kruskal-Wallis test with Dunn’s multiple comparison test compared to WT (n = 12); L159R p = 0.0076 (n = 8), T196K p > 0.9999 (12), E209K p > 0.9999 (11), E219Q p > 0.9999 (11), M303V p > 0.9999 (11), S309P p > 0.9999 (10), Y365C p > 0.9999 (12), G369D p > 0.9999 (12), I368T p = 0.6015 (12).

We next used coefficient of variation (CV) analysis to investigate whether transport of the protein to nerve terminals is impacted by these variants. Most SYT1 variants displayed a punctate distribution, indicative of efficient presynaptic targeting of the protein, and had a similar CV to the WT protein (Fig 2b). However, the L159R variant was more diffusely localised, displaying a significantly lower CV than WT (Fig 2a, b), which is in keeping with its lower nerve terminal expression (Fig 1c). We then ascertained whether SYT1 variants are correctly trafficked to synaptic vesicles, their functional site of action. SYT1 variants were tagged at their lumenal domain with pHluorin, a pH-sensitive GFP which has quenched fluorescence in acidic environments, such as the lumen of synaptic vesicles. We sequentially perfused neurons with saline, acidic, or ammonia buffer to differentiate the partitioning of SYT1 variants to distinct membrane compartments (Fig 2c). All SYT1 variants displayed a similar targeting to vesicles as the WT protein (Fig 2d). Combined, these results suggest that the majority of SYT1 variants do not significantly affect the expression or localisation of SYT1 protein within neurons; however, the L159R variant substantially reduces synaptic expression of SYT1.

**Figure 2.**
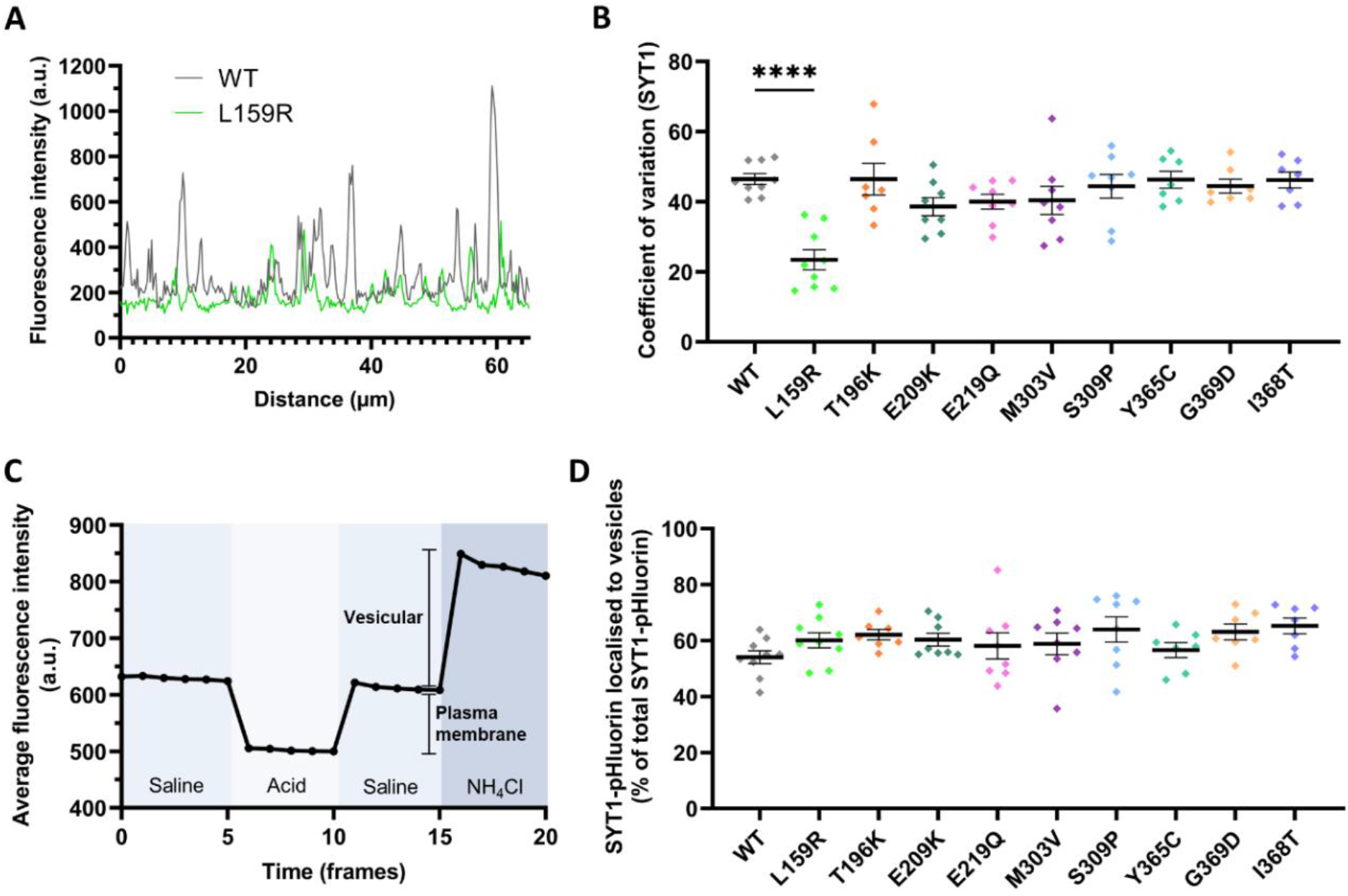
Most SYT1 variants are trafficked to and distributed within the presynaptic nerve terminal as efficiently as WT protein. Hippocampal neurons transfected with SYT1-pHluorin variants were sequentially perfused with saline, acidic, or ammonia buffers. (**A**) Representative SYT1-pHluorin fluorescence traces for CV analysis. Peaks in fluorescence intensity indicate higher densities of SYT1. High CV equates to a punctate localisation of SYT1, indicative of enrichment at presynaptic nerve terminals. (**B**) Localisation of SYT1 variants to nerve terminals expressed as CV. Data displayed as mean ± SEM, n = 7-9 (from at least 3 independent cultures). One-way ANOVA with Dunnett’s multiple comparison test compared to WT (n = 9); L159R p = <0.0001 (n = 9), T196K p >0.9999 (7), E209K p = 0.2691 (8), E219Q p = 0.4936 (8), M303V p = 0.5580 (8), S309P p = 0.9987 (8), Y365C p >0.9999 (7), G369D p = 0.9993 (7), I368T p >0.9999 (7). (**C**) Protocol and representative data for membrane partitioning assay to distinguish between plasma membrane-bound and synaptic vesicle-localised SYT1. (**D**) Vesicular localisation of SYT1 variants within the presynaptic nerve terminal compared to WT, expressed as a percentage of total SYT1-pHluorin fluorescence in the nerve terminal. Data displayed as mean ± SEM, n = 7-9 (from at least 3 independent cultures). Kruskal-Wallis test with Dunn’s multiple comparison test compared to WT (n = 9); L159R p > 0.9999 (n = 9), T196K p = 0.4231 (7), E209K p > 0.9999 (8), E219Q p > 0.9999 (8), M303V p > 0.9999 (8), S309P p = 0.1454 (8), Y365C p > 0.9999 (7), G369D p = 0.2279 (7), I368T p = 0.0904 (7).

### Effect of SYT1 variants on evoked exocytosis

Having established that most variants are unlikely to cause haploinsufficiency, we next explored whether these variants had dominant-negative impacts on SYT1 function. Firstly, we investigated whether any variants impact the total availability of synaptic vesicles for exocytosis. Neurons were stimulated with 1200 action potentials at 10 Hz in the presence of bafilomycin A1 to block vesicle reacidification. Increases in pHluorin fluorescence thus provides a measure of cumulative vesicle exocytosis. This approach was used to establish the proportion of vesicles that are mobilised by the stimulus train, which comprises the recycling pool of vesicles. The presence of SYT1 C2B domain variants M303V and S309P significantly reduced the size of the recycling pool compared to WT (Fig 3a) whereas all C2A variants (Fig 3b) and remaining C2B variants (Fig 3a) did not impact the total availability of synaptic vesicles for exocytosis.

**Figure 3.**
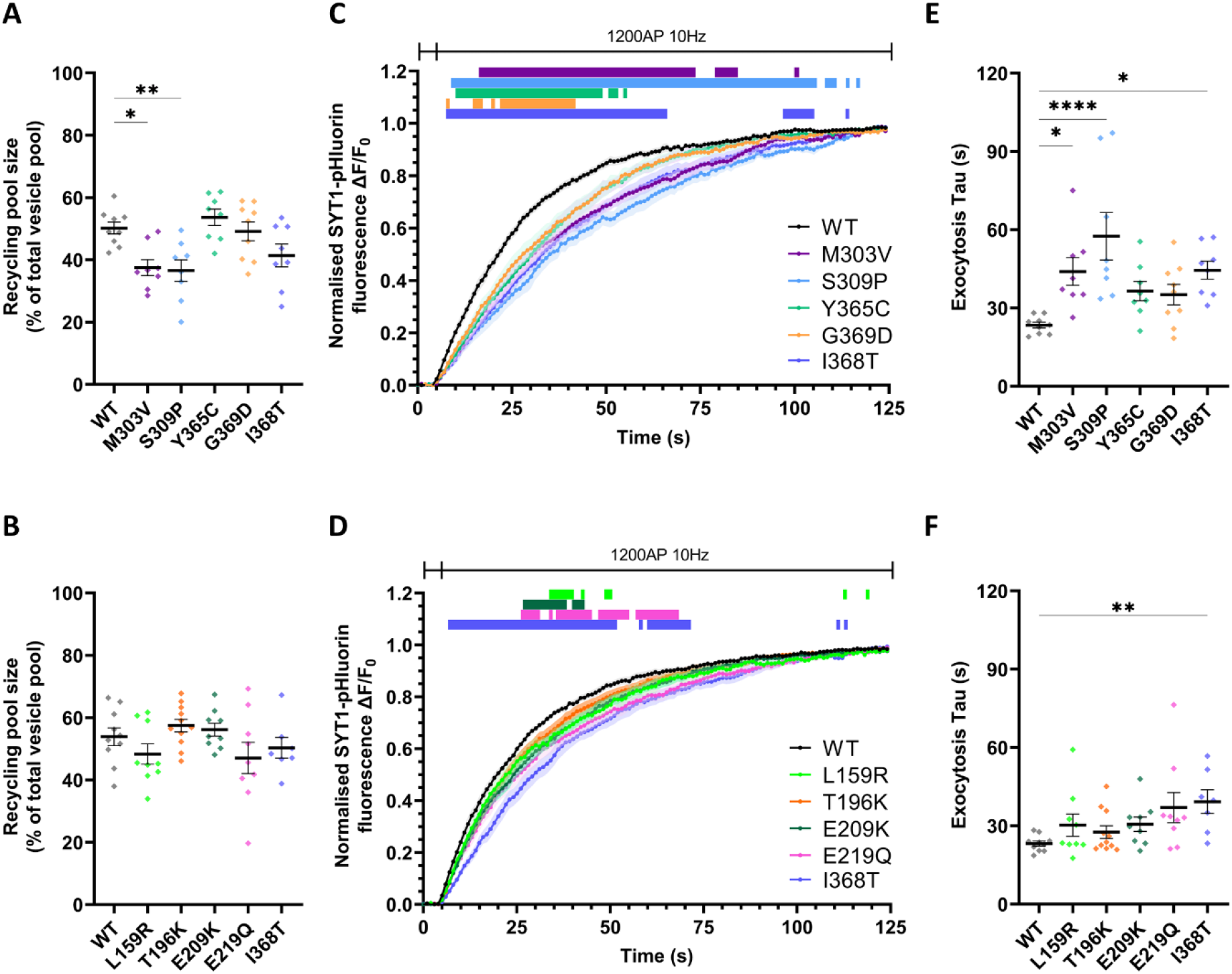
C2B and C2A domain SYT1 variants induce a dominant-negative slowing of exocytosis. Hippocampal neurons transfected with SYT1-pHluorin variants were stimulated with 1200 AP at 10 Hz in the presence of bafilomycin A1, and then perfused with ammonia buffer to reveal total SYT1-pHluorin fluorescence. (**A, B**) Synaptic vesicle recycling pool sizes in the presence of C2B (A) or C2A (B) domain variants, expressed as a percentage of the total pool of vesicles (peak SYT1-pHluorin fluorescence with ammonia buffer). One-way ANOVA with Dunnett’s multiple comparison test compared to WT; A: M303V p = 0.0143, S309P p = 0.0077, Y365C p = 0.8692, G369D p = 0.9991, I368T p = 0.1361. B: L159R p = 0.6012, T196K p = 0.8718, E209K p = 0.9826, E219Q p = 0.4059, I368T p = 0.9139. (**C, D**) Time course of mean ΔF/F_0_ of C2B (C) or C2A (D) domain SYT1-pHluorin variants normalised to peak amplitude of fluorescence change induced by stimulation. Coloured bars above graph represent time points of significant difference between the SYT1 variants and WT, analysed through repeated measures ANOVA with Dunnett’s multiple comparison test. (**E, F**) One-phase curves were fit over the entire stimulation period, from which tau constant values were obtained; tau values shown for C2B (E) or C2A (F) domain variants. One-way ANOVA with Dunnett’s multiple comparison test compared to WT for C2B variants; E: M303V p = 0.0206, S309P p = <0.0001, Y365C p = 0.2322, G369D p = 0.2976, I368T p = 0.0170. Kruskal-Wallis test with Dunn’s multiple comparison test compared to WT for C2A variants; F: L159R p = 0.8385, T196K p > 0.9999, E209K p = 0.2283, E219Q p = 0.0742, I368T p = 0.0092. All data displayed as mean ± SEM, n = 7-11 (from at least 3 independent cultures; WT n = 9 for A, C, E, n = 10 for B, D, F; I368T n = 8 for A, C, E, n = 7 for B, D, F; M303V n = 8; S309P = 8; Y365C = 8; G369D = 9; L159R = 9; T196K = 11; E209K = 9; E219Q = 9).

We then explored whether the SYT1 variants had any dominant-negative impacts on the kinetics of neurotransmitter release. The cumulative mobilisation of recycling pool vesicles was ascertained from pHluorin fluorescence time traces, normalised to the maximal fluorescence induced by stimulation (Fig 3c, d). The presence of either SYT1 C2B (Fig 3c) or C2A (Fig 3d) domain variants impaired evoked exocytosis of recycling pool vesicles in a dominant-negative manner. Notably, SYT1 variants produced graded impairments to neurotransmitter release, as evidenced by the number of time points where they differ from WT (see bars denoting significance in Fig 3c and 3d, though note that T196K is not significantly different to WT). In particular, the C2B domain variants generally had a more pronounced effect than the C2A variants (Fig 3c and 3d). This was further demonstrated through analysis of time constant (tau) values extracted from one-phase curves fit to the time courses as a measure of overall exocytic rate. Among the C2B domain variants, M303V and S309P caused a significant decrease in the overall rate of mobilisation of the recycling pool of vesicles, as was also observed for the previously-characterised I368T variant (Fig 3e). In contrast, none of the C2A domain variants had exocytic taus that significantly differed from WT (Fig 3f).

To gain a better appreciation of how SYT1 variants exert their pathogenic effects, we specifically examined synaptic vesicle exocytosis over the initial 5 seconds of stimulation when the exocytic rate is largely linear. For these analyses, we examined the rate of fusion relative to the entire pool of vesicles, to account for any changes in recycling pool size. All C2B domain SYT1 variants demonstrated a significant slowing of initial exocytic rate compared to the WT protein (Fig 4a, b). Graded effects on exocytosis kinetics were again observed across these variants; M303V and S309P variants decreased the initial rate of exocytosis to a similar extent as the I368T reference variant, whereas Y365C and G369D exhibited a less severe effect (Fig 4a, b). C2A domain variants all induced a modest reduction in initial exocytic rate but this effect was not statistically significant (Fig 4c, d). Together, these findings suggest that SYT1 variants in both the C2A and C2B domains impair synaptic vesicle exocytosis; however, they do so to varying degrees, with C2B domain variants having substantially greater impacts on evoked exocytosis compared to C2A domain variants.

**Figure 4.**
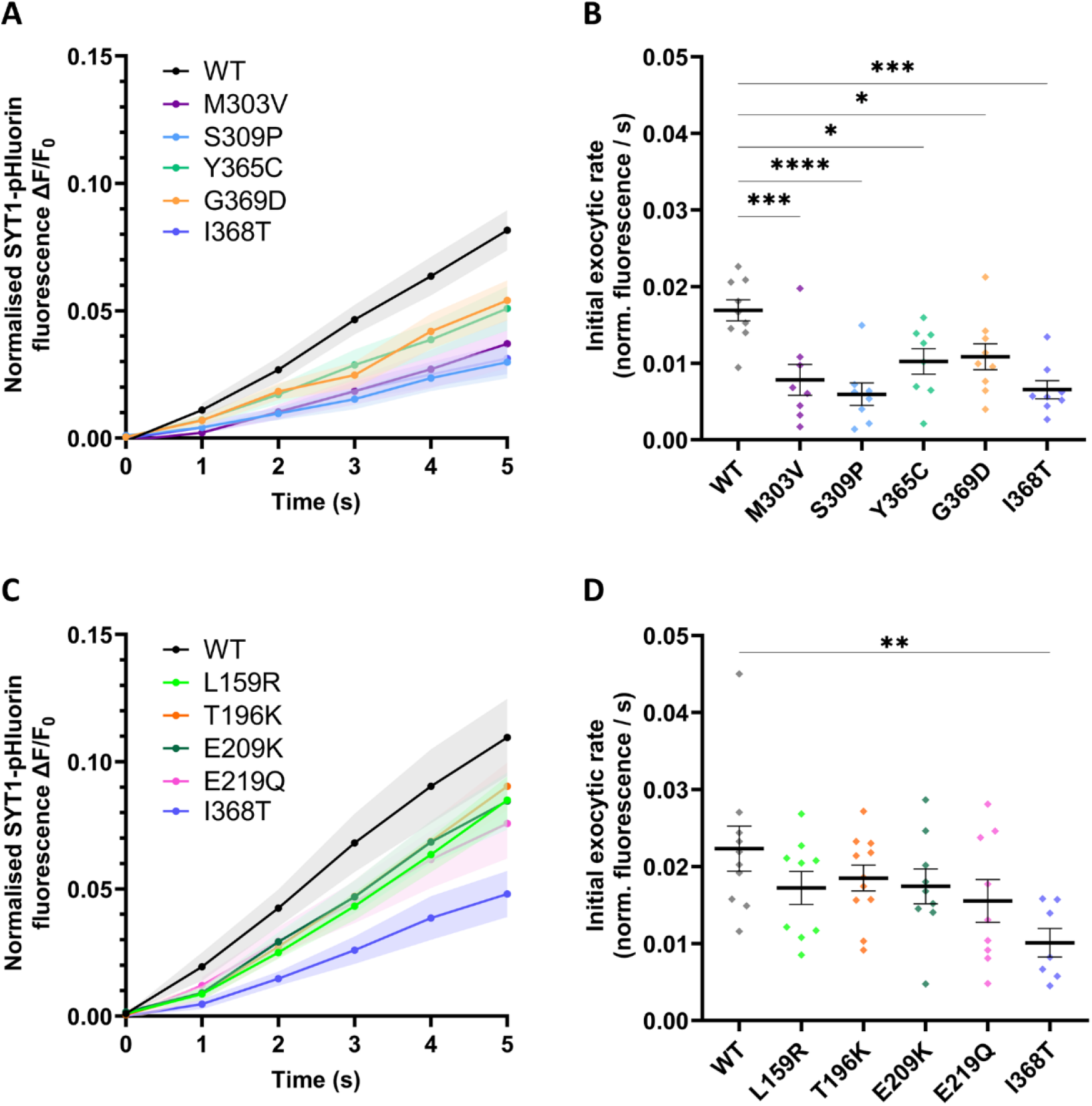
Impact of SYT1 variants on initial exocytic rate. Hippocampal neurons transfected with SYT1-pHluorin variants were stimulated with 1200 AP at 10 Hz in the presence of bafilomycin A1 and perfused with ammonia buffer to reveal total SYT1-pHluorin fluorescence. (**A, C**) Time course (first 5 seconds of stimulation) of mean ΔF/F_0_ of C2B (A) or C2A (C) domain SYT1-pHluorin variants normalised to maximum fluorescence induced by ammonia buffer. (**B, D**) Initial exocytic rates for C2B (B) or C2A (D) domain variants over the first 5 seconds of stimulation. One-way ANOVA with Dunnett’s multiple comparison test compared to WT; B: M303V p = 0.0009, S309P p = <0.0001, Y365C p = 0.0196, G369D p = 0.0318, I368T p = 0.0001. D: L159R p = 0.4016, T196K p = 0.6250, E209K p = 0.4403, E219Q p = 0.1596, I368T p = 0.0048. Data displayed as mean ± SEM, n = 7-11 (from at least 3 independent cultures; WT n = 9 for A, B, n = 10 for C, D; I368T n = 8 for A, B, n = 7 for C, D; M303V n = 8; S309P = 8; Y365C = 8; G369D = 9; L159R = 9; T196K = 11; E209K = 9; E219Q = 9).

### Associations between presynaptic functional deficits and phenotyping measures for participants with SYT1 variants

We next tested whether the degree of exocytic impairments induced by SYT1 variants is associated with variation in the neurodevelopmental phenotypes of individuals harbouring these variants. Analyses were focused on neurodevelopmental characteristics which we have previously reported to differ between SYT1 participants and comparison participants with other monogenic intellectual disabilities (Melland et al., 2022): communication and motor impairments through subscales of the Vineland Adaptive Behaviour Scales (VABS), cerebral visual impairments (CVI), movement disorders, and self-injurious behaviour from the Developmental Behaviour Checklist (DBC). We explored correlations between quantitative phenotyping data for each of the SYT1 variants examined in this paper (except L159R due to data unavailability) and three different measures of presynaptic functional impact thatreflect different aspects of vesicle fusion and neurotransmitter release: initial exocytic rate within the first 5 seconds, overall exocytic rate (tau, equating to rate of mobilisation of the recycling pool), and percentage of vesicles fused by 200 AP (timepoint where SYT1 variants induced their strongest impacts, Supp Fig 1). Table 1 shows permutation correlation test results for associations between phenotypic measures and metrics of synaptic vesicle exocytosis dysfunction. Strong correlations were found between the percentage of vesicles exocytosed by 200 AP and age-standardised scores on both the Vineland motor and the Vineland communication sub-scales (Table 1, Fig 5a and b), such that a greater functional deficit correlated with more severe impairments in adaptive functioning. Strong correlations were also found between the Vineland motor sub-scale and both the initial exocytic rate and exocytic tau, where a lower initial exocytic rate or a slower tau, respectively, correlated with more severe motor deficits (Table 1, Fig 5a and b). In contrast, no metric of presynaptic dysfunction significantly correlated with the number of movement disorder types reported, DBC self-injury sub-scale, or CVI score (Table 1, Fig 5b, Supp Fig 2-4).

**Figure 5.**
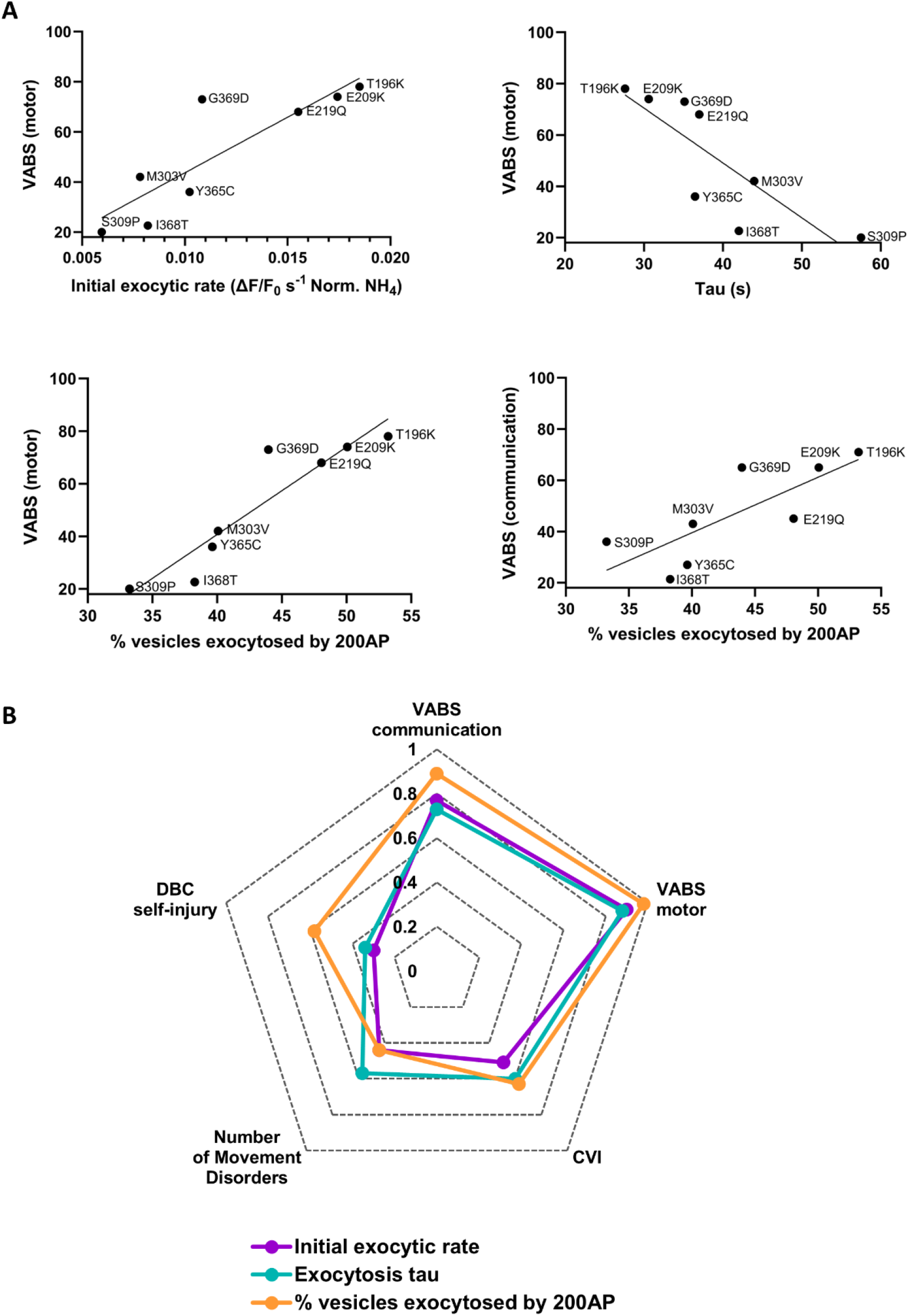
Exocytic defects induced by SYT1 variants are strongly correlated with motor and communication phenotypes of SYT1-associated neurodevelopmental disorder. Quantitative phenotypes exhibited by individuals harbouring SYT1 variants were each assessed for correlation with 3 separate functional measures of exocytosis kinetics – initial exocytic rate (over first 5 seconds of 10Hz stimulation), overall exocytic rate (tau), and percentage of vesicles fused by 200AP (Supp Fig 1). Phenotyping measures include motor and communication subscales of the Vineland Adaptive Behaviour Scale (VABS), number of movement disorders, self-injury scores from the Developmental Behaviour Checklist (DBC), and cerebral visual impairment (CVI). (**A**) Scatter plots of significant correlations between exocytic measures in the presence of SYT1 variants and quantitative neurodevelopmental phenotypic measures of individuals with SYT1 variants. (**B**) Radar plot of correlations between exocytic measures in the presence of SYT1 variants and neurodevelopmental phenotypic measures of individuals with SYT1 variants. Spearman’s r coefficients are presented with directionality removed for ease of interpretation.

**Table 1.** Correlations between exocytosis metrics and participant phenotyping measures.

| | Initial exocytic rate<br>(norm. ($\Delta F/F_0$)/s) | | Exocytosis<br>tau (s) | | % vesicles fused by<br>200AP | |
| --- | --- | --- | --- | --- | --- | --- |
|  | *r* | *p* | *r* | *p* | *r* | *p* |
| VABS communication | 0.77 | 0.028 | -0.73 | 0.042 | 0.89 | 0.004* |
| VABS motor | 0.90 | 0.002* | -0.88 | 0.005* | 0.98 | < 0.001* |
| CVI | -0.51 | 0.241 | 0.60 | 0.164 | -0.63 | 0.129 |
| Number of Movement Disorders | -0.44 | 0.252 | 0.57 | 0.138 | -0.44 | 0.249 |
| DBC Self-Injury | -0.30 | 0.552 | 0.34 | 0.506 | -0.58 | 0.222 |

Asterisks indicate significant results after corrections. Significance threshold at $p \leq 0.019$ to account for multiple comparisons. VABS = Vineland Adaptive Behaviour Scale; DBC = Developmental Behaviour Checklist; CVI = cerebral visual impairment.

## Discussion

Here, we have established the pathogenicity of newly identified SYT1 C2B variants and, for the first time, variants in the C2A domain of the protein. We show that variants in disparate regions of SYT1 share a common underlying mechanism of pathogenicity, causing a graded dominant-negative impairment in synaptic vesicle exocytosis. Importantly, we have also explored genotype-function-phenotype relationships for these variants, demonstrating that the extent of presynaptic impairment correlates with motor and communication difficulties in individuals harbouring these variants (Fig 6). This establishes impairment of evoked neurotransmitter release as a likely candidate mechanism constraining the development of cognitive and motor abilities, leading to core features of SYT1-associated neurodevelopmental disorder.

**Figure 6.**
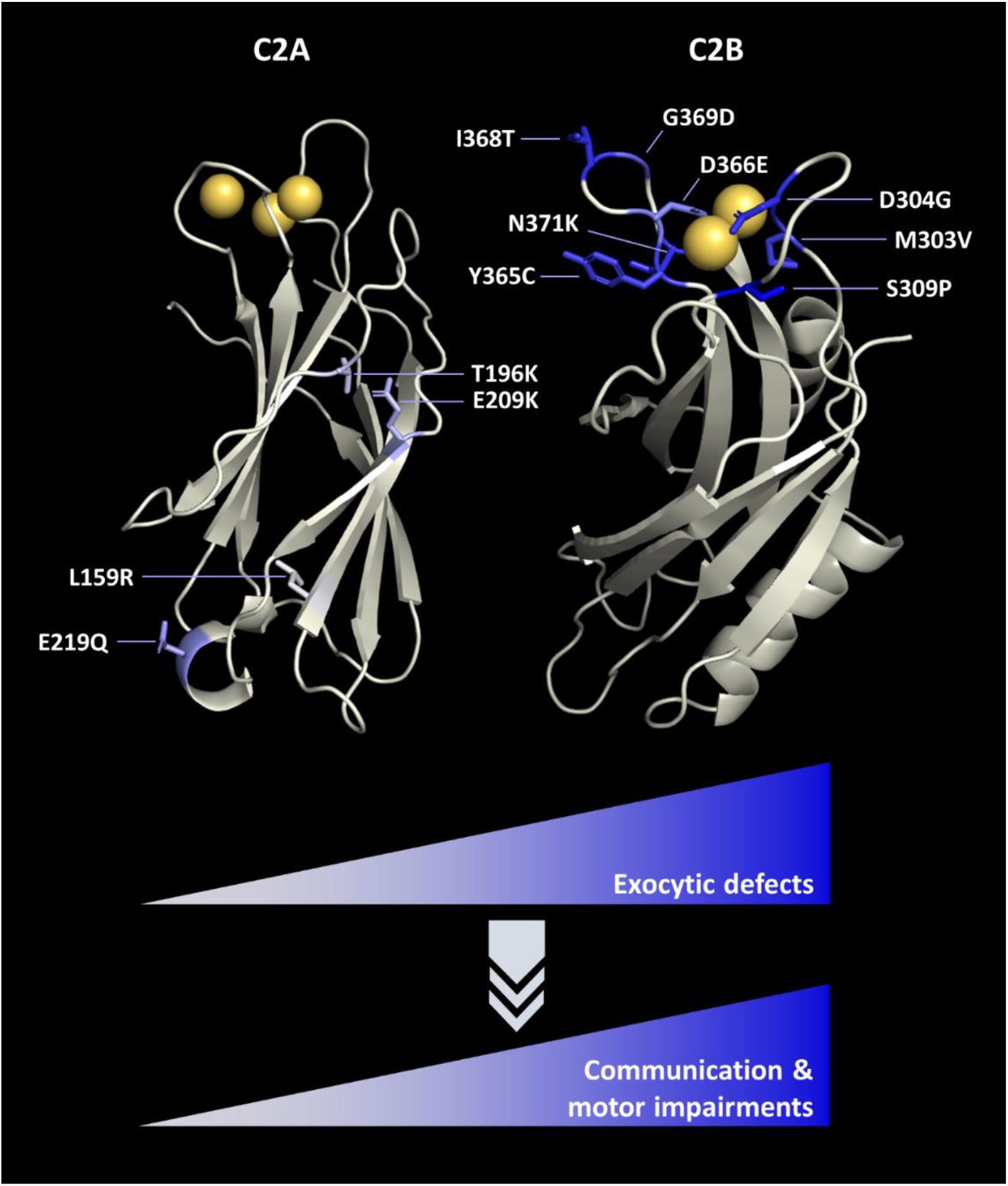
Genotype-function-phenotype relationship in SYT1-associated neurodevelopmental disorder. SYT1 variants in both the C2A domain (top left) and C2B domain (top right) cause graded, dominant-negative impairments to evoked exocytosis (this paper, (Baker et al., 2018), colour of each residue equates to relative impact to exocytosis). The degree of impaired exocytosis correlates with motor and communication difficulties in individuals harbouring these variants, and is dependent on the nature of the amino acid substitution and its location within SYT1. SYT1 variants that perturb protein stability tend to present with stronger adaptive functions and fewer neurological symptoms, which may be due to either reduced expression or mitigation of dominant-negative effects.

We demonstrate that four recently identified SYT1 C2B domain variants impair synaptic vesicle exocytosis in a dominant-negative manner, a pathogenic mechanism previously established for other SYT1 C2B variants (Baker et al., 2015; Baker et al., 2018). Evoked exocytosis was reduced over both shorter and prolonged periods of stimulation in the presence of the C2B variants, though the degree of impact was variable. The position of variants within the Ca^2+^-binding pocket of the C2B domain may contribute to this variability. Variants located in Ca^2+^-binding loop 1 (M303V, S309P (this study) and D304G (Baker et al., 2018)) cause severe exocytic defects, whereas some loop 3 variants, including Y365C, G369D (this study) and D366E (Baker et al., 2018), induced milder exocytic impairments. However, this is not axiomatic: N371K and I368T (this study and (Baker et al., 2018)), which induce comparatively severe exocytic deficits, also reside in loop 3. Moreover, alternative variants at a recurrent locus can have differential effects. M303K was previously shown to impair the stability and expression of SYT1 (Baker et al., 2018), an effect not recapitulated by M303V in this current study. Intriguingly, these variants are also associated with different neurodevelopmental characteristics (Melland et al., 2022; Baker et al., 2018). Therefore, the functional severity, or indeed mechanism of action, of variants cannot be simply predicted by residue location alone.

Importantly, we provide the first functional evidence for the pathogenicity of C2A domain variants identified in individuals with neurodevelopmental delay. C2A domain variants act in a similar fashion to C2B variants, impairing synaptic vesicle exocytosis in a dominant-negative manner. However, C2A domain variants had comparatively modest impacts and did not significantly reduce the initial exocytic rate; instead, significant divergence from WT SYT1 only emerged with extended stimulation. Overall, the severity of exocytic impairment induced by all four C2A variants was milder than the comparison C2B I368T variant. It is noteworthy, however, that E219Q had the greatest impact among the C2A variants, as evidenced by the time course of cumulative exocytosis. Correspondingly, the individual harbouring this variant also presented with more severe impairments in adaptive behaviour than the other C2A variants (Melland et al., 2022). The severity of this particular C2A variant was not predictable from molecular modelling, highlighting the power of functional interrogation of SYT1 variants to not only establish their mechanism of pathogenicity, but also for predicting neurodevelopmental phenotypes in SYT1-associated neurodevelopmental disorder.

These findings suggest that the impact of SYT1 variants in either C2 domain converge on a similar synaptic phenotype, with C2B domain variants having, on average, a greater magnitude of dominant-negative effect on exocytosis compared to C2A domain variants. This observation is in line with the differential proposed roles of the two domains, whereby the C2B domain is essential for triggering evoked exocytosis, while the C2A domain plays an auxiliary role (Striegel et al., 2012; Gruget et al., 2020; Lee et al., 2013). However, there is a notable confound to the simple comparison between variants in C2A and C2B domains: in contrast to the C2B variants explored here, all C2A variants are found outside the Ca^2+^-binding loops (Fig 6). In fact, all investigated C2A variants occur in regions of currently unknown significance to SYT1 function. For most C2A variants (except L159R, as detailed below), molecular dynamics simulations and modelling did not indicate any major impacts on domain structure or function or obvious mechanism of pathogenicity (Melland et al., 2022). Hence, it is perhaps unsurprising that these variants have milder impacts on exocytosis kinetics overall compared to their C2B domain counterparts, most of which involve substitutions to residues of the critical Ca^2+^-binding loops. It remains to be seen whether variants affecting Ca^2+^-binding residues of the C2A domain would result in mild or more severe exocytic defects (and corresponding clinical impacts).

We also explored alternative mechanisms of pathogenicity beyond dominant-negative effects, by examining how SY1 variants impact protein expression and transport. In keeping with observations from C2B variants studied to date (Baker et al., 2015; Baker et al., 2018; Melland et al., 2023), most of the investigated SYT1 variants across the C2A and C2B domains had no impact on nerve terminal expression or trafficking of SYT1, indicating that disruption to protein stability and expression are not common consequences of SYT1 missense variants.

There are, however, exceptions to this rule - the C2A domain variant L159R impaired SYT1 expression and targeting to nerve terminals. L159 lies in a beta-strand and its sidechain faces the hydrophobic interior of the C2A beta-sandwich structure. Substituting a hydrophobic sidechain (leucine) for a positively charged sidechain (arginine) was predicted via molecular dynamic simulations to introduce structural instability to multiple regions of the C2A domain (Melland et al., 2022). The aforementioned M303K C2B domain variant, which also increased structural instability, was similarly expressed at lower levels (Baker et al., 2018). Interestingly, from qualitative reports (Cafiero et al., 2015; Melland et al., 2022; Baker et al., 2018) both variants are associated with a relatively mild neurodevelopmental phenotype. It is therefore possible that these two variants confer haploinsufficiency and/or have a limited ability to exert deleterious dominant negative effects in neurons. The latter argument is supported by the current evidence that, despite its impaired expression, some copies of SYT1 L159R can traffic to vesicles and can exert dominant-negative slowing of exocytosis, albeit mild. Thus, amino acid substitutions to diverse regions of the protein can have major impacts to stability and expression, likely dependent on the nature of the variant. Together, these findings indicate that SYT1 variants may have heterogeneous molecular and cellular effects which are not easily predicted by variant location, though dominant-negative impairment of exocytosis remains the most frequent pathogenic mechanism.

This study has specifically investigated missense variants that were selected because they 1) fall within different domains of the protein, 2) reside in different regions of each domain, and/or 3) are predicted to have diverse impacts on protein structure. Further insights will be gained by investigating a greater number of variants, increasing power for correlational analyses. For example, it would be important to explore whether a recently identified C2A variant (V184A) identified in an individual with a severe, early-lethal neurodevelopmental disorder (Huang et al., 2023), causes severe impairment to evoked neurotransmitter release. Moreover, we have only assessed the impacts of these variants on evoked exocytosis; previously investigated SYT1 C2B variants slow evoked exocytosis without having dominant-negative impacts on the additional functions of SYT1 as a clamp for spontaneous neurotransmitter release and a modulator of endocytosis (Melland et al., 2023). It therefore remains possible that the newly-identified SYT1 variants may additionally impact these other presynaptic processes.

Nevertheless, the strong correlations observed between exocytic impairments and severity of adaptive impairments support disruption to evoked neurotransmitter release as a primary pathogenic mechanism underlying SYT1-associated neurodevelopmental disorder. We have previously observed that three individuals harbouring the SYT1 C2B variant D366E presented with milder motor delay than those harbouring other SYT1 variants, and did not exhibit the early-onset movement disorders seen in other members of the cohort; this variant also induced the mildest impact on evoked exocytosis among those variants examined (Baker et al., 2018). However, these observations were based on non-standardised, qualitative reporting of clinical phenotypes and only a single variant associated with milder exocytic defects and clinical presentations. In the present study, we aggregated presynaptic function and quantitative phenotyping data for 8 SYT1 variants across the two C2 domains, which demonstrated a significant correlation between standardised scores of motor and communication skills and metrics of exocytic efficacy. This represents the first evidence for a unifying functional impairment that underlies and drives these adaptive impairments.

We have focussed our study on a limited range of phenotypes that we have previously detailed to be prevalent in SYT1-associated neurodevelopmental disorder, and for which quantitative measures were available. The observed function-phenotype correlations did not extend to all phenotype measures, with no significant correlations between impairments in exocytosis observed *in vitro* and the number of movement disorders, cerebral visual impairment scores or self-harm items of the DBC. Methodological limitations include the lack of sensitivity of carer-report measures for these phenotypic dimensions, and limited power to detect significant associations in a small group size, especially for complex behavioural characteristics which will be subject to modifying influences and changes over time. We also note that for most variants phenotyping data was only available for a single individual, though for recurrent variants phenotypic variation has been shown to be remarkably consistent (Melland et al., 2022). Thus, the absence of significant correlations should be interpreted with caution and does not preclude mechanistic relationships for these symptom domains. However, if the observed contrasts in function-phenotype associations are later confirmed, this could reflect diverse mechanisms contributing to different aspects of the condition.

Our results inform knowledge of the mechanism of dysfunction caused by SYT1 variants at the level of single synapses, whilst the complex SYT1-associated neurodevelopmental phenotypes implicate widespread brain networks maturing across development. What then might link impaired exocytosis to motor and communication deficits? There are several possible explanations. Reduced neurotransmitter release may influence synaptogenesis and/or embryonic formation of local and distributed neuronal circuitry (Kochubey et al., 2016; Cooper & Gillespie, 2011). However, major disruptions to SYT1 function do not substantially alter neuronal pathfinding, or synapse number and morphology in multiple cellular and *in vivo* mammalian and invertebrate model systems (Liu et al., 2009; Chicka et al., 2008; Geppert et al., 1994; Fernandez-Chacon et al., 2002; Imig et al., 2014; Yoshihara & Littleton, 2002; Huson et al., 2020), though SYT1 has been shown to positively influence axonal branching in an avian model (Greif et al., 2013). Notably, even in the absence of evoked neurotransmitter release, such as in prenatal SNAP-25 knockout mice, axonal growth and thalamocortical development is largely unperturbed (Washbourne et al., 2002).

A more likely possibility is that dampening of evoked synchronous release reduces the fidelity of neurotransmission. Impaired evoked neurotransmitter release would be expected to have an ongoing impact on synaptic strength and plasticity, blunting processes such as long-term potentiation, impairing learning and memory and thereby leading to constraints on the acquisition and optimisation of motor and cognitive skills (Xu et al., 2012). The impacts that this has on discrete circuits and thus behaviours would depend on the relative contribution of SYT1 to these. There are up to 17 members of the synaptotagmin family, and in particular SYT2, SYT7 and SYT9 replace SYT1 as the major sensors for fusion in a number of central neuronal cell types (Xu et al., 2007; Chen et al., 2017; Bacaj et al., 2013; Weber et al., 2014). Reliance on SYT1, or sensitivity to impaired evoked synchronous release, therefore likely varies between specific cell types, circuits, brain regions and developmental time-windows (Banerjee et al., 2020; Marqueze et al., 1995; Ullrich et al., 1994; Turecek & Regehr, 2019). Therefore, impaired SYT1 function is expected to lead to diverse impacts to neural networks and circuits causing varied behavioural impairments that may change across the lifespan.

Further *in vivo* experimental evidence is required to resolve the multi-level mechanisms bridging between parameters of presynaptic function and the emergent properties of motor control and cognitive development. Developing *in vivo* animal models harbouring neurodevelopmental disorder-associated SYT1 variants will be key to understanding how these affect whole-brain, network-based systems. These models could subsequently be used to explore whether elements of SYT1-associated neurodevelopmental disorder can be ameliorated. With the finding that perturbed exocytosis is common to both C2A and C2B variants, and correlates with disease severity, this emerges as a core tractable target to investigate for therapeutic amelioration of this disorder.

## Data availability

Raw data and images are available upon request from the corresponding author.

## Abbreviations

4-AP: 4-aminopyridine
AP: action potential
CNQX: cyanquixaline, 6-cyano-7- nitroquinoxaline-2,3-dione
CV: coefficient of variation
CVI: cerebral visual impairment
DBC: Developmental Behaviour Checklist
DL-AP5: DL-2-amino-5-phosphonopentanoic acid
EGFP: enhanced green fluorescent protein
MES: 2-(N-morpholino)ethanesulfonic acid
ROI: region of interest
SYT1: synaptotagmin-1
VABS: Vineland Adaptive Behaviour Scales
WT: wild-type

## Acknowledgements & Funding

We are grateful to the study participants and parents / carers who contributed to this paper. The Florey Institute of Neuroscience and Mental Health acknowledges the strong support from the Victorian Government and in particular the funding from the Operational Infrastructure Support Grant. For the purpose of open access, the author has applied CC BY public copyright licence to any Author Accepted Manuscript version arising from this submission. This work was supported by National Health and Medical Research Council (NHMRC) Ideas Grant (2003710) to SG, and UK Medical Research Council (G101400) and Great Ormond Street Hospital Children’s Charity to KB. PYP acknowledges the support of an Australian Government Research Training Program Scholarship.

## Author Contribution

Conceptualisation: S.L.G, K.B. and P.Y.P.; Data Curation: P.Y.P., L.E.B., J.E.; Formal Analysis: P.Y.P., L.E.B., N.S., R.A.J.; Funding Acquisition: S.L.G. K.B.; Investigation: P.Y.P., L.E.B., N.S., M.A.A., J.E., R.A.J.; Visualisation: P.Y.P. and L.E.B.; Writing-original draft: P.Y.P. (lead), S.L.G.; Writing-review and editing: P.Y.P., S.L.G., K.B., L.E.B., H.M. All other authors reviewed and approved this manuscript.

