## Supplementary Material for "Adaptive functions correlate with evoked neurotransmitter release in SYT1-associated neurodevelopmental disorder"

### Materials

SYT1-pHluorin (a pH-sensitive variant of GFP) was provided by Prof. V. Haucke (Leibniz Institute of Molecular Pharmacology, Berlin, Germany). QuikChange II Site-Directed Mutagenesis kit was from Agilent Technologies (Santa Clara, CA, USA). Thermo Plasmid Maxiprep Kit, Neurobasal media, B-27 supplement, penicillin/streptomycin, Minimal Essential Medium (MEM), Dulbecco's MEM/F-12, Lipofectamine 2000, goat anti-chicken IgY (H+L) Alexa Fluor 488 and donkey anti-rabbit IgG (H+L) DyLight 550 were obtained from ThermoFisher Scientific (Scoresby, Australia). Rabbit anti-SYT1 was from Synaptic Systems (Göttingen, Germany). Chicken anti-GFP was from Abcam (Cambridge, UK). 6-cyano-7- nitroquinoxaline-2,3-dione (CNQX) was from ENZO Life Sciences (Lausen, Switzerland). DL-2-Amino-5-phosphonopentanoic acid (DL-AP5) was from Caymen Chemical (Ann Arbor, MI, USA). Bafilomycin A1 was from Toronto Research Chemicals (Toronto, Canada). All other reagents were obtained from Sigma-Aldrich (Castle Hill, Australia).

### Methods

#### Site-directed mutagenesis

I368T SYT1-pHluorin was previously generated as described (Baker et al., 2015). Mutagenesis of SYT1-pHluorin was performed using the QuickChange II kit (Agilent) to introduce the human SYT1 missense variants (L159R, T196K, E209K, E219Q, M303V, S309P, Y365C, G369D) into the homologous position in rat SYT1 using primers listed in Table 1. Human sequence numbering is used throughout for clarity. Successful mutagenesis of plasmid DNA was confirmed through Sanger sequencing performed by the Australian Genome Research Facility, Melbourne, Australia. Transfection-grade plasmids were produced using the Thermo Plasmid Maxiprep Kit as per the manufacturer's instructions.

**Table 1. Primers used for site-directed mutagenesis of SYT1-pH variants.** Mutated bases in bold and underlined

| SYT1 Variant (Human) | Primer | Sequence (5'-3') |
|---|---|---|
| L159R | Forward<br>Reverse | CCAGAATAACCAGCG**G**TTGGTGGGAATCATC<br>GATGATTCCCACCAA**C**CGCTGGTTATTCTGG |
| T196K | Forward<br>Reverse | CCTGACAAAAAGAAGAAATTTGAGA**AG**AAAGTCCACCGGAAAACCC<br>GGGTTTTCCGGTGGACTTT**CT**TCTCAAATTTCTTCTTTTTGTCAGG |
| E209K | Forward<br>Reverse | CCCTCAATCCAGTCTTCAAT**A**AACAATTTACTTTCAAGGTACCCTA<br>CCGAGTAGGGTACCTTGAAAGTAAATTGTT**T**ATTGAAGACTGGATT |
| E219Q | Forward<br>Reverse | CAAGGTACCCTACTCG**C**AATTAGGTGGCAAAACCC<br>GGGTTTTGCCACCTAATT**G**CGAGTAGGGTACCTTG |

| | | |
|---|---|---|
| M303V | Forward<br>Reverse | CCTGAAGAAG**G**TGGATGTGGGTGGCTTATCTG<br>CAGATAAGCCACCCACATCCA**C**CTTCTTCAGG |
| S309P | Forward<br>Reverse | GATGGATGTGGGTGGCTTA**C**CTGATCCCTACG<br>CGTAGGGATCAG**G**TAAGCCACCCACATCCATC |
| Y365C | Forward<br>Reverse | GGTGGTAACTGTTTTGGACT**G**TGACAAGATTGGCAAGAACG<br>CGTTCTTGCCAATCTTGTCA**C**AGTCCAAAACAGTTACCACC |
| G369D | Forward<br>Reverse | GGACTATGACAAGATTG**A**CAAGAACGACGCGATCGGC<br>GCCGATCGCGTCGTTCTTG**T**CAATCTTGTCATAGTCC |

**Primary hippocampal neuronal culture**

All procedures were approved by the Florey Animal Ethics Committee and performed in accordance with the guidelines of the National Health and Medical Research Council Code of Practice for the Care and Use of Animals for Experimental Purposes in Australia. C57BL/6J mouse in-house colonies were maintained in a temperature controlled (≈ 21°C) room and group housed in individually ventilated cages on a 12 hour light-dark cycle (lights on 07:00–19:00) with food and water available ad libitum. Animals were time mated overnight and visualisation of a vaginal plug on the following morning was considered as embryonic day (E) 0.5.

Dissociated primary hippocampal-enriched neuronal cultures were prepared from hippocampi dissected from at least three E16.5-18.5 C57BL/6J mouse embryos of both sexes. Hippocampi were incubated in 10 units/mL papain for 20 minutes at 37°C; papain was removed, and the hippocampal tissue was mechanically disaggregated in pre-equilibrated DMEM/F-12 media supplemented with 10% foetal bovine serum and 1% penicillin/streptomycin. The cell suspension was centrifuged at 363 g for 5 minutes, and the hippocampal cell pellet was resuspended in fresh, pre-equilibrated Neurobasal (NB) media supplemented with 1% penicillin-streptomycin, 0.5mM L-glutamine and 1x B-27 (henceforth termed “enriched NB”). Harvested cells were subsequently plated onto poly-D-lysine and laminin-coated coverslips at a density of 35,000 cells/coverslip for 24-well plates (on 13mm coverslips) and 60,000 cells/coverslip for 6-well plates (on 25mm coverslips), after which they were incubated for an hour at 37°C, 5% $CO_2$ before supplementation with pre-equilibrated enriched NB media. At 3-4 days *in vitro* (i.e. DIV3-4), cultures were further supplemented with 1µM 1-β-D-arabinofuranosylcytosine to inhibit glial proliferation.

**Transfection**

At DIV6-7, the hippocampal neuronal cultures were co-transfected with SYT1-pHluorin variants in the presence of either pcDNA empty vector for fixed cell imaging, or mCherry for live cell imaging assays. Lipofectamine 2000 (1 uL/well for 24-well plates, 2 uL/well for 6-well plates) was incubated in MEM (125 uL/well for 24-well plates, 250 uL/well for 6-well plates) for 5 minutes at room temperature. Plasmid DNA (0.5 ug/construct/well for 24-well plates, 1 ug/construct/well for 6-well plates) in MEM media (125 uL/well for 24-well plates, 250 uL/well for 6-well plates) was combined with an equal volume of lipofectamine mixture and incubated at room temperature for 20 minutes. Enriched NB media was collected from cells and retained, and cells were transfected in pre-equilibrated MEM for 2 hours at 37°C, 5% $CO_2$. The cells were gently washed with MEM and the conditioned enriched NB media replaced, and cells maintained at 37°C, 5% $CO_2$ until use.

#### Fixation & Immunocytochemistry

At DIV13 transfected primary hippocampal neuronal cultures were first washed with saline imaging buffer (in mM: 136 NaCl, 2.5 KCl, 2 CaCl2, 1.3 MgCl2, 10 glucose, 10 HEPES, pH 7.4) and then fixed in 4% paraformaldehyde in phosphate-buffered saline (PBS) for 20 minutes, incubated at room temperature in 50mM $NH_4Cl$ in PBS for 10 minutes, washed with PBS and permeabilised with 0.1% v/v Triton x100, 1% v/v bovine serum albumin (BSA) in PBS for 5 minutes. Cells were then washed with PBS and blocked with 1% BSA in PBS for one hour. Cells were subsequently immunolabelled with primary antibodies in 1% BSA in PBS (1:2000 chicken anti-GFP, Abcam, Cat#: ab13970; 1:100 rabbit anti-SYT1, SYSY, Cat#: 105102) in a dark humidified chamber for 1.5 - 2 hours, washed extensively with PBS, then immunolabelled in the dark with secondary antibodies in 1% BSA in PBS (1:500 488 Alexafluor goat anti-chicken, Cat#: A11039; 1:200 550 DyLight donkey anti-rabbit, Cat#: SA5-10039, both ThermoFisher Scientific) over 1 hour. Finally, all cells were labelled with 4',6-diamidino-2- phenylindole (DAPI) for maximum of 5 minutes. Cells were washed 3 times with PBS after each incubation period to remove excess antibodies. Coverslips were then rinsed twice in fresh double-distilled $H_20$, then mounted on glass slides using Dako Mounting Medium.

#### Fluorescence Imaging

Fixed, immunolabelled cells or live neuronal cultures were imaged on a Zeiss Axio Observer 7 inverted epifluorescence microscope with a Colibri 7 LED light source. Live neuronal cultures on 25mm coverslips were mounted in a Warner imaging chamber with embedded parallel platinum wires (RC-21BRFS) before being placed on the stage of the microscope. All neurons were visualised through a Zeiss EC Plan-Neofluar 40x/1.30 oil-immersion objective and a Zeiss Axiocam 506 mono camera. 14-bit images were captured with 2x2 binning yielding a pixel size of 0.227 µm.

SYT1-pHluorin was visualised with a 488 nm excitation wavelength through a single bandpass GFP filter set (excitation 470/40, emission 525/50, beam splitter 495 nm). SYT1 and cell soma were visualised through AF555 and DAPI channels respectively, with excitation wavelengths of 553 and 353 nm respectively through a quadruple bandpass 90 HE filter set (excitation 385/20, 469/28, 555/20, 632/24 and emission 425/30, 514/30, 591/25, 709/100, beam splitter 405/493/575/653 nm). mCherry was visualised with a 545 nm excitation wavelength through a single bandpass DsRed filter set (excitation 550/24, emission 605/70, beam splitter 570 nm).

For live imaging, all buffers were warmed to 37°C via a Warner inline heater, and neurons were stimulated via a D343 Dual Stimulator module employed in a D330 MultiStim System (SDR

Scientific Pty Ltd, Chatswood, Australia). Automated triggering of electrical stimulation and variable image acquisition rates were implemented using the Zen Blue Experiment Designer module.

For the membrane partitioning assay, live hippocampal neuronal cultures were perfused sequentially with saline imaging buffer (as described above), 20mM 2-ethanesulfonic acid (MES) buffer (in mM: 136 NaCl, 2.5 KCl, 2 $CaCl_2$, 1.3 $MgCl_2$, 10 glucose, 20 MES, pH 5.5), and 50mM $NH_4Cl$ alkaline imaging buffer (in mM: 86 NaCl, 2.5 KCl, 2 $CaCl_2$, 1.3 $MgCl_2$, 10 glucose, 10 HEPES, 50 $NH_4Cl$, pH 7.4. For the exocytosis assay, live cultures were perfused with saline imaging buffer (as described above, supplemented with 10 µM CNQX, 50 µM DL-AP5 and 1 µM bafilomycin A1) at 37°C. Neurons were stimulated with 1,200 action potentials at 10Hz (35mA, 1ms pulse width) for 2 min. Neurons were then perfused with an alkaline $NH_4Cl$ buffer (as described above) to reveal the total pHluorin fluorescence. Images were captured at 1s intervals.

**Imaging analysis**

Images were analysed using the FIJI 1.52n distribution of ImageJ and the Time Series Analyzer V3 plugin. Nerve terminal expression analysis of fixed cell images were carried out by first selecting 8x8 pixel regions of interest (ROIs) over transfected and non-transfected nerve terminals and background regions in the GFP channel. SYT1 expression level in transfected neurons was calculated using Microsoft Excel as fold-change over the non-transfected endogenous SYT1 expression level by dividing the average fluorescence intensity of transfected puncta by that of non-transfected puncta after subtracting background intensity.

To correct for slight XY-plane drift over the time series, stack registration was performed by rigid body alignment to the first frame of the experiment using the MBGReg ImageJ plugin (Donal Stewart, Edinburgh, UK), a custom variant of the FIJI StackReg plugin. For live cell imaging analysis, 10x10 pixel ROIs were selected over transfected nerve terminals as well as background. To remove the component of fluorescence decay contributed by the photobleaching of background autofluorescence in time series images from evoked exocytosis assays, the Bleach Correction Image J plugin was used (version: CorrectBleach_-2.0.3-SNAPSHOT.jar, https://github.com/fiji/CorrectBleach) (Miura, 2020). Photobleaching over the full length of the experiment was corrected using a single exponential function fitted to the fluorescence decay of background ROIs over the duration of the experiment prior to $NH_4Cl$ buffer superfusion. After bleach-correcting against the selected set of background ROIs, transfected ROIs were laid over the image and synaptic fluorescence intensities measured. Each transfected ROI subsequently underwent a rigorous screening process with strict inclusion/exclusion criteria. Only ROIs that increased fluorescence from baseline upon both electrical stimulation and $NH_4Cl$ buffer superfusion were included in subsequent analysis, and coverslips with greater than 20 responsive ROIs were included in the final data.

For exocytosis assays, fluorescence responses of each ROI were calculated in Microsoft Excel as $\Delta F/F_0$ (change in fluorescence from baseline). For each frame of the time series, $\Delta F/F_0$ fluorescence values were averaged for all ROIs in a field of view. The average fluorescence intensity traces were subsequently normalised through one of two methods; Fluorescence intensities were either normalised to the peak fluorescence at the end of stimulation, or to the peak fluorescence exhibited in response to the $NH_4Cl$ alkaline buffer.

**Statistical analyses of functional data**

Statistical analyses of SYT1 variant functional data were performed using GraphPad Prism Version 9.4.0 and Microsoft® Excel software. Each data set was tested for normality using the Shapiro-Wilk test and statistical analyses were performed accordingly. Comparison between each SYT1 variant protein and the wild-type (WT) protein with regards to nerve terminal localisation, vesicle recycling pool size, and exocytosis parameters of initial exocytic rate for both C2A and C2B variants, and overall exocytic rate (tau) for C2B variants, were performed through one-way ANOVA with Dunnett's multiple comparison test after confirming normality of the data. Comparison between each SYT1 variant protein and the wild-type (WT) protein with regards to nerve terminal expression, vesicular localisation and tau for C2A variants were performed through a non-parametric Kruskal-Wallis test with Dunn's multiple comparisons due to non-normality of the data. Time courses of change in fluorescence were analysed via repeated measures mixed model two-way ANOVA with Dunnett's multiple comparison test, to compare each SYT1 variant to the WT control at every individual time point. For all analyses, $p < 0.05$ was considered statistically significant where the degree of significance is represented by: * $p < 0.05$, ** $p < 0.01$, *** $p < 0.001$, **** $p < 0.0001$. All data are displayed as mean ± SEM, where individual data points are shown as diamonds. 'n' refers to an individual field of view from an independent coverslip. Sample numbers are indicated in the graphs as single datapoints, and in the figure legends. No data that met inclusion criterion has been excluded from this study. Experimenters were blinded to the SYT1-pHluorin variant for image acquisition and analysis for fixed assays, and at the image analysis stage for live imaging. No strategy for randomization of samples has been followed, and sample sizes were not chosen based on pre-specified effect sizes. Instead, several independent experiments were performed across multiple independent cultures as indicated in figure legends, with all experiments repeated across at least 3 independent cultures (and each culture comprising at least 3 embryos). To minimise confounds, controls (WT) were performed in each batch of experiments alongside SYT1 variants, and the well-characterised recurrent C2B variant I368T was included as a reference variant in each assay.

**Statistical analyses of functional and clinical data correlations**

Permutation Spearman's correlation tests were performed to examine associations between functional activity (i.e. the synaptic vesicle exocytosis kinetics in vitro, of neurons transfected with SYT1-pHluorin variants) and clinical phenotypes of patients with the equivalent SYT1 variants. The SYT1 variants included in this analysis, based on available functional and phenotypic data, are the following: T196K, E209K, E219Q, M303V, S309P, Y365C, G369D, and I368T. To account for multiple comparisons, the false discovery rate was controlled using the Benjamini-Hochberg procedure, with $q < 0.05$. Permutation-based partial (Spearman's) correlation tests were used to measure correlations between functional activity and vision problems while controlling for age. (For other phenotypic measures, bivariate correlation tests were appropriate as the variables represented standardised scores accounting for age). Prior to conducting analyses, S309P was identified as an outlier, using Cook's distance, for the following correlation pairs: Tau and adaptive behaviour, Tau and communication problems, and Tau and motor problems. Removing S309P did not however impact the analysis outcomes as the r coefficients before and after were highly similar. In addition, the data was missing for behavioural and emotional problems for S309P. As such, the results shown included all the available dataset.

**Table 2. SYT1-associated Neurodevelopmental Disorder participant characteristics**

| | N | Mean (SD) | Range |
|---|---|---|---|
| Sex | 7 female<br>7 male | - | |
| Age[a]<br>*Years* | 14 | 9.5 (6.1) | 3.2 – 25.8 |
| Global adaptive function<br>*Vineland Adaptive Behaviour Composite* | 12 | 44.8 (18) | 20 - 74 |
| Behavioural and emotional difficulties<br>*DBC total problems T-score* | 11 | 56.6 (8.2) | 40 - 68 |

[a]Age at time of questionnaire completion (n=13) or clinical data collection (n=1)

**Table 3. SYT1-associated Neurodevelopmental Disorder participant clinical phenotypes**

| Clinical feature<br>HPO Term Identifier[a] | Data available (n) | Frequency of Feature (n) | Frequency of Feature (%) | Subtype (n) |
|---|---|---|---|---|
| Delayed speech and language development<br>HP: 0000750 | 14 | 14 | 100 | Mild = using words and phrases (5); moderate = using single words only (2); severe = not using any words (4); unable to classify as under age 5 years or insufficient information (3) |
| Abnormal eye physiology<br>HP: 0012373 | 14 | 12 | 86 | Strabismus (7); nystagmus (4); hypermetropia (1); visual impairment unspecified (1) |
| Motor delay<br>HP: 0001270 | 14 | 14 | 100 | Mild = walked by 3 years (5); moderate = walked by 5 years (2); severe = walked after 5 years or nonambulatory over the age of 5 (5); unable to classify because nonambulatory under the age of 5 years (2) |
| Neonatal hypotonia<br>HP: 0001319 | 14 | 11 | 79 | - |
| Abnormality of movement<br>HP: 0100022 | 14 | 11 | 79 | Dystonia (5); chorea (7); dyskinesia (3); ataxia (5); myoclonus (3); tremor (3); stereotypies (5); Tourette syndrome (1) |
| Sleep disturbance<br>HP: 0002360 | 13 | 9 | 69 | - |
| Abdominal symptom<br>HP: 0011458 | 14 | 8 | 57 | Feeding difficulties (6); gastroesophageal reflux (2); drooling (1); constipation (2); urinary infection (2); pancreatitis with pseudocysts (1) |
| Self-injurious behaviour<br>HP: 0100716 | 14 | 11 | 79 | Finger biting or chewing (7); head banging (2); skin picking (1); hair pulling (1); other or unspecified (2) |
| Seizure<br>HP: 0001250 | 14 | 4 | 29 | - |

| | | | | |
|---|---|---|---|---|
| MRI abnormality<br>HP: 0012639 | 13 | 3 | 23 | - |
| Abnormality of the respiratory system<br>HP: 0002086 | 13 | 2 | 15 | Hyperventilation with cyanosis (1); autonomic dysfunction with hypotension (1) |
| Phenotypic abnormality<br>HP: 0000118 | 13 | 4 | 31 | Undescended testicle (1); dermoid cyst (1); talipes (2) |
| Abnormality of prenatal development or birth<br>HP: 0001197 | 14 | 2 | 14 | Mild prematurity (1); neonatal resuscitation (2) |

[a]Human Phenotype Ontology (https://hpo.jax.org/app/).

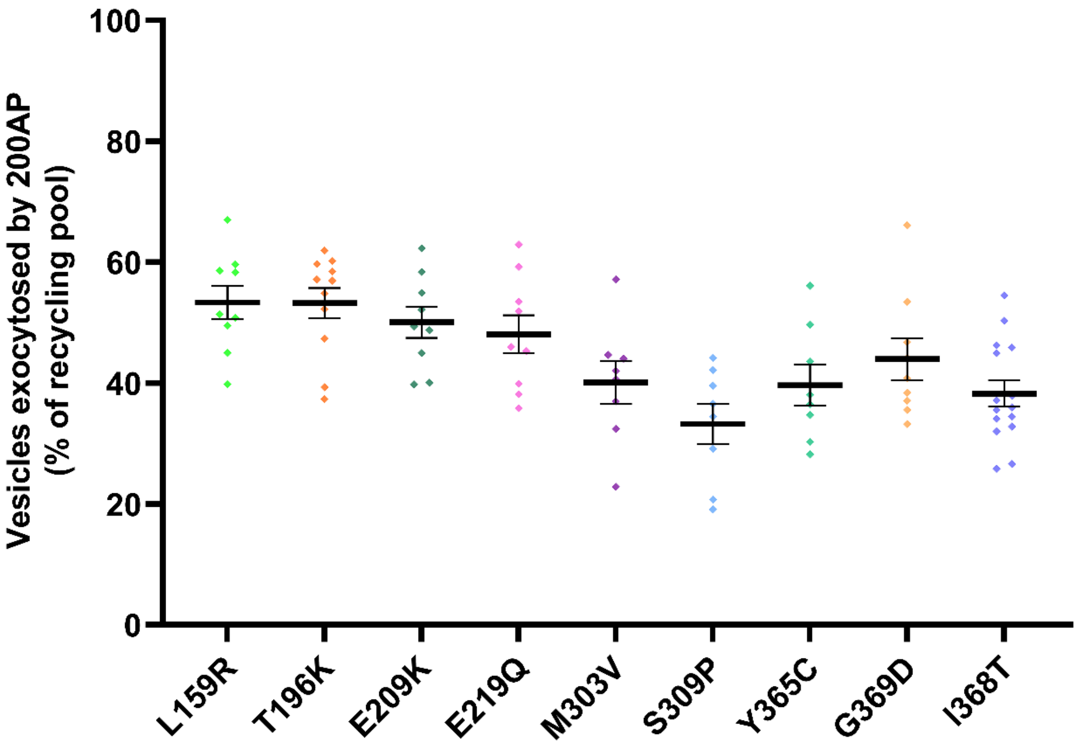


**Supplemental Figure 1. The number of synaptic vesicles exocytosed cumulatively by 200AP stimulation.** The percentage of synaptic vesicles cumulatively exocytosed following 200AP (i.e. after 20 seconds of stimulation) stimulation, as a proportion of the total recycling pool of vesicles, was determined for all SYT1 variants. Data displayed is mean +/- SEM.

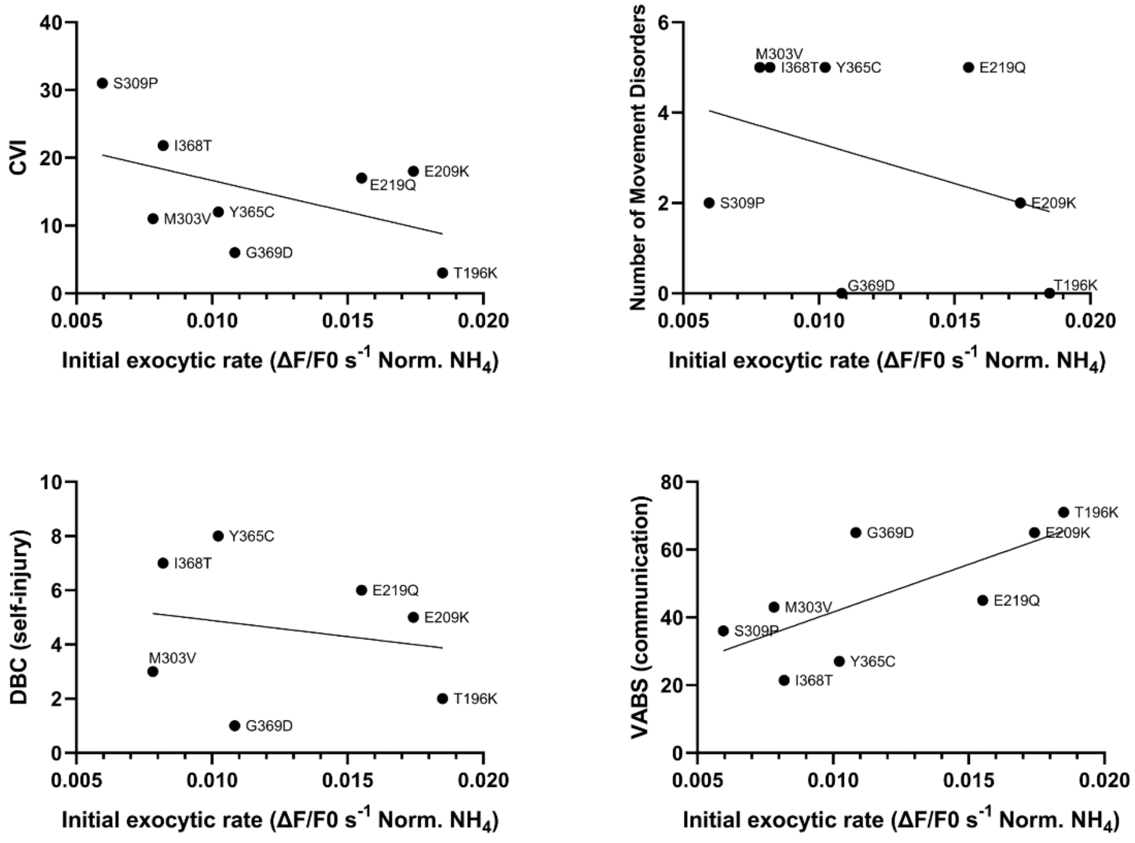


**Supplemental Figure 2. Scatter plots of non-significant correlations between initial exocytic rate in the presence of SYT1 variants and clinical measures of individuals with SYT1 variants.** Quantitative phenotypes exhibited by individuals harbouring SYT1 variants were each correlated with the functional measure of initial exocytic rate over the first 5 seconds of 10Hz stimulation. Clinical measures include the communication subscale of the Vineland Adaptive Behaviour Scale (VABS), number of movement disorders, self-injury scores from Developmental Behaviour Checklist (DBC), and cerebral visual impairment (CVI). Correlation values can be found in Table 1 of the main text.

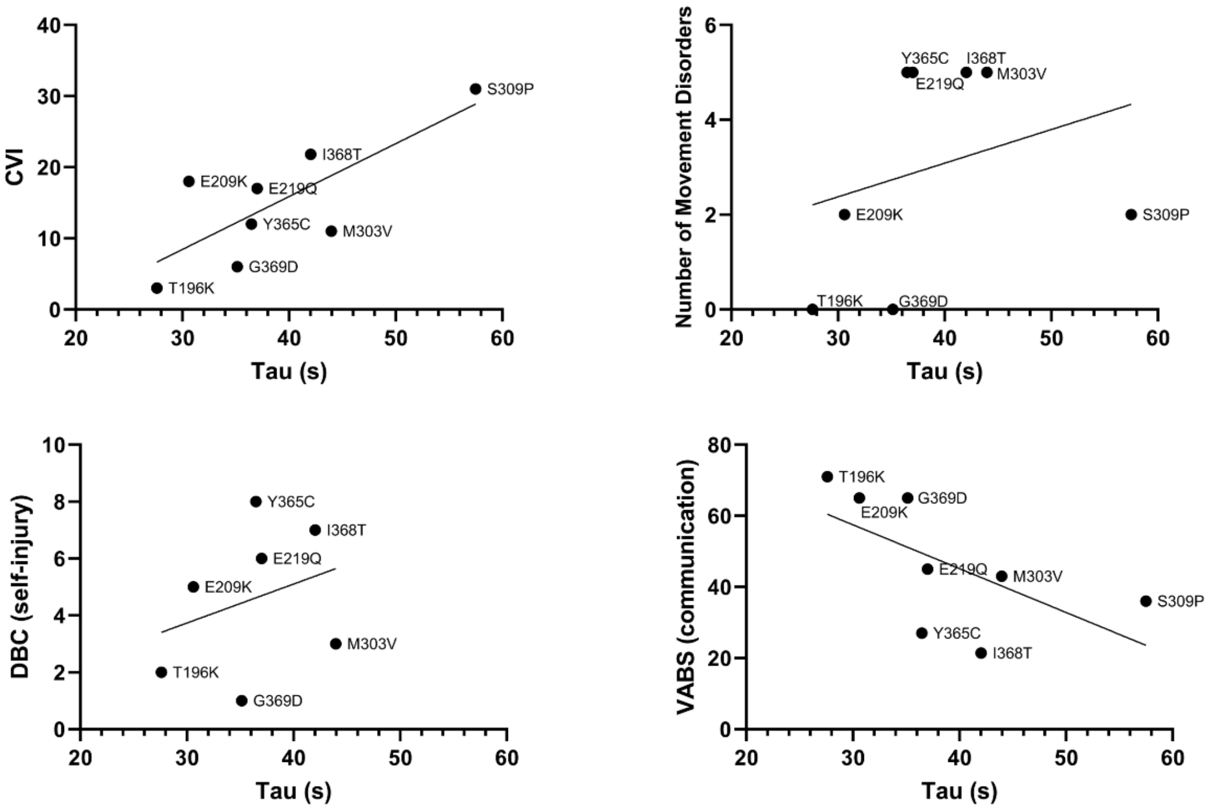


**Supplemental Figure 3. Scatter plots of non-significant correlations between tau constant values in the presence of SYT1 variants and clinical measures of individuals with SYT1 variants.** Quantitative phenotypes exhibited by individuals harbouring SYT1 variants were each correlated with the functional measure for overall exocytic rate (tau). Clinical measures include the communication subscale of the Vineland Adaptive Behaviour Scale (VABS), number of movement disorders, self-injury scores from Developmental Behaviour Checklist (DBC), and cerebral visual impairment (CVI). Correlation values can be found in Table 1 of the main text.

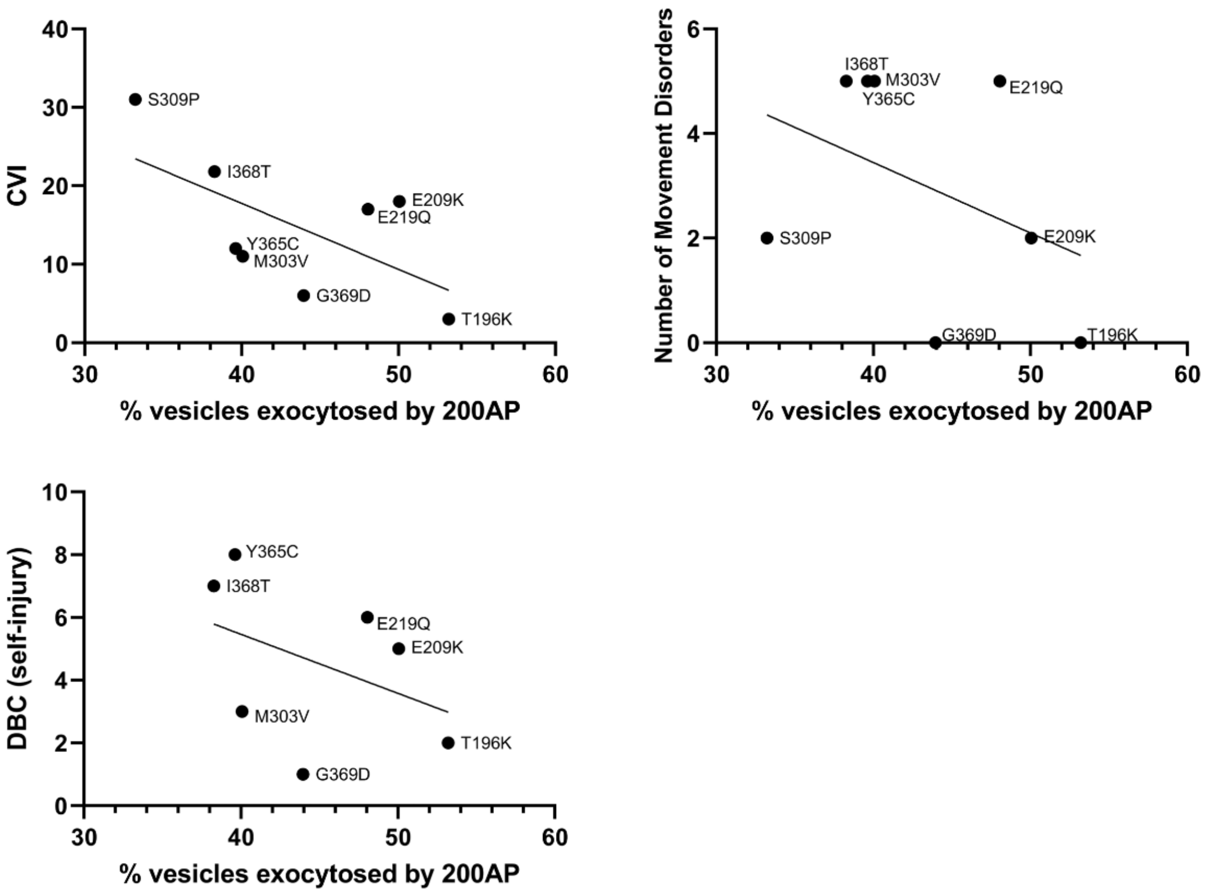


**Supplemental Figure 4. Scatter plots of non-significant correlations between the percentage of versicles fused by 200AP in the presence of SYT1 variants and clinical measures of individuals with SYT1 variants.** Quantitative phenotypes exhibited by individuals harbouring SYT1 variants were each correlated with the functional measure of percentage of vesicles fused by 200AP. Clinical measures include number of movement disorders, self-injury scores from Developmental Behaviour Checklist (DBC), and cerebral visual impairment (CVI). Correlation values can be found in Table 1 of the main text.
